# GABA_A_ receptor mapping in human using non-invasive electrophysiology

**DOI:** 10.1101/2020.05.11.087726

**Authors:** Alexander D Shaw, Hannah L Chandler, Khalid Hamandi, Suresh D Muthukumaraswamy, Alexander Hammers, Krish D Singh

## Abstract

The non-invasive study of cortical oscillations provides a window onto neuronal processing. Temporal correlation of these oscillations between distinct anatomical regions is considered a marker of functional connectedness. As the most abundant inhibitory neurotransmitter in the mammalian brain, γ-aminobutyric acid (GABA) is thought to play a crucial role in shaping the frequency and amplitude of oscillations, which thereby suggests a role for GABA in shaping the topography of functional activity and connectivity. This study explored the effects of pharmacologically blocking the reuptake of GABA (increasing local concentrations) through oral administration of the GABA transporter 1 (GAT1) blocker tiagabine (15 mg). We show that the spatial distribution of tiagabine-induced activity changes, across the brain, corresponds to group-average flumazenil PET maps of GABA_A_ receptor distribution.

In a placebo-controlled crossover design, we collected resting magnetoencephalography (MEG) recordings from 15 healthy male individuals prior to, and at 1-, 3- and 5- hours post, administration of tiagabine and placebo pill. Using leakage-corrected amplitude envelope correlations (AECs), we quantified the functional connectivity in discrete frequency bands across the whole brain, using the 90-region Automatic Anatomical Labelling atlas (AAL90), as well as quantifying the average oscillatory activity across the brain.

Analysis of variance in connectivity using a drug-by-session (2×4) design revealed interaction effects, accompanied by main effects of drug and session. Post-hoc permutation testing of each post-drug recording against the respective pre-drug baseline revealed consistent reductions of a bilateral occipital network spanning theta, alpha and beta frequencies, and across 1- 3- and 5- hour recordings following tiagabine, but not placebo.

The same analysis applied to activity, across the brain, also revealed a significant interaction, with post-hoc permutation testing demonstrating significant increases in activity across frontal regions, coupled with reductions in activity in posterior regions, across the delta, theta, alpha and beta frequency bands.

Crucially, we show that the spatial distribution of tiagabine-induced changes in oscillatory activity overlap significantly with group-averaged maps of the estimated distribution of GABA_A_ receptors, derived from scaled flumazenil volume-of-distribution (FMZ-V_T_) PET, hence demonstrating a possible mechanistic link between GABA availability, GABA_A_ receptor distribution, and low-frequency network oscillations. We therefore propose that electrophysiologically-derived maps of oscillatory connectivity and activity can be used as sensitive, time-resolved, and targeted receptor-mapping tools for pharmacological imaging at the group level, providing direct measures of target engagement and pharmacodynamics.

## Introduction

There is a pressing need to develop time-resolved non-invasive markers of pharmacological effects within the brain, that can be used to both drive our understanding of brain function but also demonstrate the action and temporal dynamics of novel pharmacological agents. This could help reduce the very significant costs associated with the development of new drugs, by helping to demonstrate direct target engagement, and therefore accelerate the development of novel treatments for diseases that have significant societal burdens. One strong candidate for such a marker are neural oscillations, either modulated by task or characterised at rest. Unlike indirect methods, such as functional magnetic resonance imaging (fMRI), neural oscillations measured using non-invasive methods, such as magnetoencephalography (MEG), arise directly from the synchronous oscillatory-activation of post-synaptic currents in ensembles of principal cells in cortex (Brunel & Wang, 2003; Buzsáki et al., 2004; Wang, 2010), predominantly those in superficial layers (Buzsáki et al., 2012; Xing et al., 2012). Converging evidence from experimental (Kramer et al., 2008; Whittington et al., 1995) and computational (Brunel & Wang, 2003; Pinotsis et al., 2013; Shaw et al., 2017) studies suggests that the attributes of these oscillations – the peak frequency and amplitude – are determined by more fundamental, unobserved, neurophysiological processes. For example, the peak frequency of oscillations within the alpha range (8 – 13 Hz) may be a mechanistic consequence of synchronised firing (action potentials) at the same rate (Halgren et al., 2017; Lorincz et al., 2009), while the peak frequency of oscillations across many frequency bands has been linked with inhibitory neurotransmission (Buzsáki et al., 2004; Hall et al., 2011; Roopun et al., 2006; Whittington et al., 2000) mediated by γ-aminobutyric acid (GABA).

The link between GABA and macroscopic oscillations observed in MEG is likely mediated by the functional inhibitory role of GABA receptors, which are broadly categorised as fast, ionotropic GABA_A_ and slower G-protein coupled GABA_B_, although there exist subtypes of each (Belelli et al., 2009; Vargas, 2018; Whiting, 2003). The currents mediated by both of these receptor types have been implicated in oscillation generation through control of recurrent excitation-inhibition (Atallah & Scanziani, 2009; Brunel & Wang, 2003; Galarreta & Hestrin, 1998; Kohl & Paulsen, 2010; Krupa et al., 2014), whereby the GABAergically-mediated inhibitory currents exert temporal control over the firing and activity of excitatory principal cells (Bartos et al., 2007; Whittington et al., 2000). In the case of higher-frequency oscillations, including beta (13 - 30 Hz) and gamma (30+ Hz), the peak frequency may be dependent on the balance of excitatory and inhibitory currents (Brunel & Wang, 2003; Kujala et al., 2015; Shaw et al., 2017), implicating excitatory glutamatergic receptor types, such as α-amino-3-hydroxy-5-methyl-4-isoxazolepropionic acid (AMPA) and N-methyl-D-aspartate (NMDA).

Pharmaco-MEG studies provide a framework for examining the effects that pharmacological manipulation of neurotransmitter or neuromodulator systems has on oscillatory activity. To date, studies of GABAergic drugs have primarily examined changes in task-induced fluctuations in oscillation metrics, usually confined to the analysis of one particular sensory system. For example, in the motor cortex, benzodiazepines, such as diazepam, modulate the amplitude of movement-related beta oscillations (Hall et al., 2011; Jensen et al., 2005). In visual cortex, propofol, an agonist of GABA_A_ receptors, increased the amplitude of gamma oscillations and concurrently suppressed the amplitude of alpha oscillations (Saxena et al., 2013). Tiagabine, a GABA re-uptake inhibitor, reduced the frequency of stimulus-induced gamma oscillations (Magazzini et al., 2016). Alcohol, which has a complex binding profile including benzodiazepine-mediated GABA_A_ effects, decreased the frequency and increased the amplitude of gamma in the visual cortex (Campbell et al., 2014). For a summary of other GABAergic pharmaco-MEG studies, see (Muthukumaraswamy, 2014; Nutt et al., 2015) These results clearly demonstrate a role for GABA in shaping oscillatory responses, however they are confined to observations of task-related, local dynamics.

Non-task induced recordings of oscillatory activity, at ‘resting state’, focus on both activity and correlation of fluctuations in different brain regions, within specific frequency bands. This ‘functional connectivity’ approach rests on the assumption that those discrete regions of the brain whose band-limited amplitude envelopes are correlated, are functionally connected. Since GABA plays a role in shaping the oscillations of local (within-region), task-induced dynamics, it is expected that it also shapes the topography of the networks formed by these regions interacting.

No pharmaco-M/EEG studies have examined GABAergic manipulation of functional connectivity across the brain, although studies examining other compounds have. Antagonism of NMDA receptors by ketamine decreased alpha and beta networks spanning motor, parietal and occipital regions (Muthukumaraswamy et al., 2015) while antagonism of AMPA receptors by perampanel increased alpha and beta in similar posterior visual and parietal regions (Routley et al., 2017). Comparison of the spatial distribution of these reductions with recent maps of receptor distributions (Zilles & Palomero-Gallagher, 2017) revealed that these areas also have higher-than-average densities of NMDA and AMPA receptors in superficial cortical layers, as well as higher-than-average densities of GABA_A_ and GABA_B_ receptors (where the average is calculated over the 44 cortex-wide regions tested) (Zilles & Palomero-Gallagher, 2017).

In the present study, we examined the effect of pharmacologically blocking the reuptake of GABA using tiagabine, a GABA-transporter 1 (GAT1) blocker. We anticipated that, at rest, increased GABA availability would lead to increased activation of GABAergic receptors and thereby GABAergic inhibition, which would have a macroscopic effect of reducing functional connectivity strengths across frequency bands.

Having identified regions demonstrating tiagabine-induced band-limited changes in oscillatory activity, we next examined the degree to which these changes overlap with the spatial distribution of GABA_A_ receptors in a group-average map of scaled flumazenil volume-of-distribution (FMZ-V_T_) PET. Here, we expected that brain regions showing the highest volume-of-distribution of FMZ-V_T_ (interpreted as the density of GABA_A_ receptors) would also demonstrate the biggest drug-induced changes in oscillatory activity.

## Materials and Methods

### MEG Methods

#### Sample

Fifteen healthy individuals (1 female, 14 male) aged between 20 – 32 (mean 25.5) completed a single-blind, placebo-controlled crossover study of 15mg oral tiagabine (Gabitril) and placebo. Study days were separated by at least a week, with each day consisting of 4 MEG scans; an initial ‘pre-drug’ recording, followed by post-ingestion scans at 1, 3 and 5 hours. MEG data from the active task protocols in this study, including a full overview of the design, has been published previously (Magazzini et al., 2016; Muthukumaraswamy et al., 2013). All study participants gave informed consent and the study procedures were approved by the UK National Research Ethics Service (South East Wales). Volunteers were excluded if they reported personal history of psychiatric or neurological diagnosis, current recreational or prescription drug use, or impaired liver function, indicated by conventional liver function test.

#### Data Acquisition

Whole head MEG recordings were obtained using a CTF275 radial gradiometer system, with participants seated upright. Task-free, ‘resting’ recordings of 10 minutes were obtained at 1200 Hz and analysed as synthetic third-order gradiometers (Vrba, 2001). Participants were fitted with three electromagnetic head coils (nasion and bilateral pre-auriculars), which remained fixed throughout the study day, and were localized relative to the MEG system immediately before and after recording. All study participants had a T1-weighted MRI (1mm isotropic) available for subsequent source-space analysis. Co-registration was achieved by placement of fiduciary markers at fixed distances from anatomical landmarks easily identifiable on an anatomical MRI (tragus, eye centre). Offline, recordings were downsampled to 600 Hz and segmented into 2 s epochs. Each epoch was visually inspected for gross artifact (e.g. movement, clenching, eye blinks) and removed from the analysis if present.

#### Resting-state analysis pipeline

The present analysis sought to compute the functional connectivity between a set of spatially resolved brain loci. We chose to compute this using the amplitude envelope correlation (AEC) metric, which is interpretable, robust and repeatable (Colclough et al., 2016). This coupling was computed separately for 7 distinct frequency bands using conventional definitions; delta (1 – 4 Hz), theta (4 – 8 Hz), alpha (8 – 13 Hz), beta (13 - 30 Hz) and 3 gamma windows (40 – 60, 60 – 80 and 80 – 100 Hz). Figure 1 shows a graphical representation of the analysis pipeline, which is detailed below.

**Figure 1.**
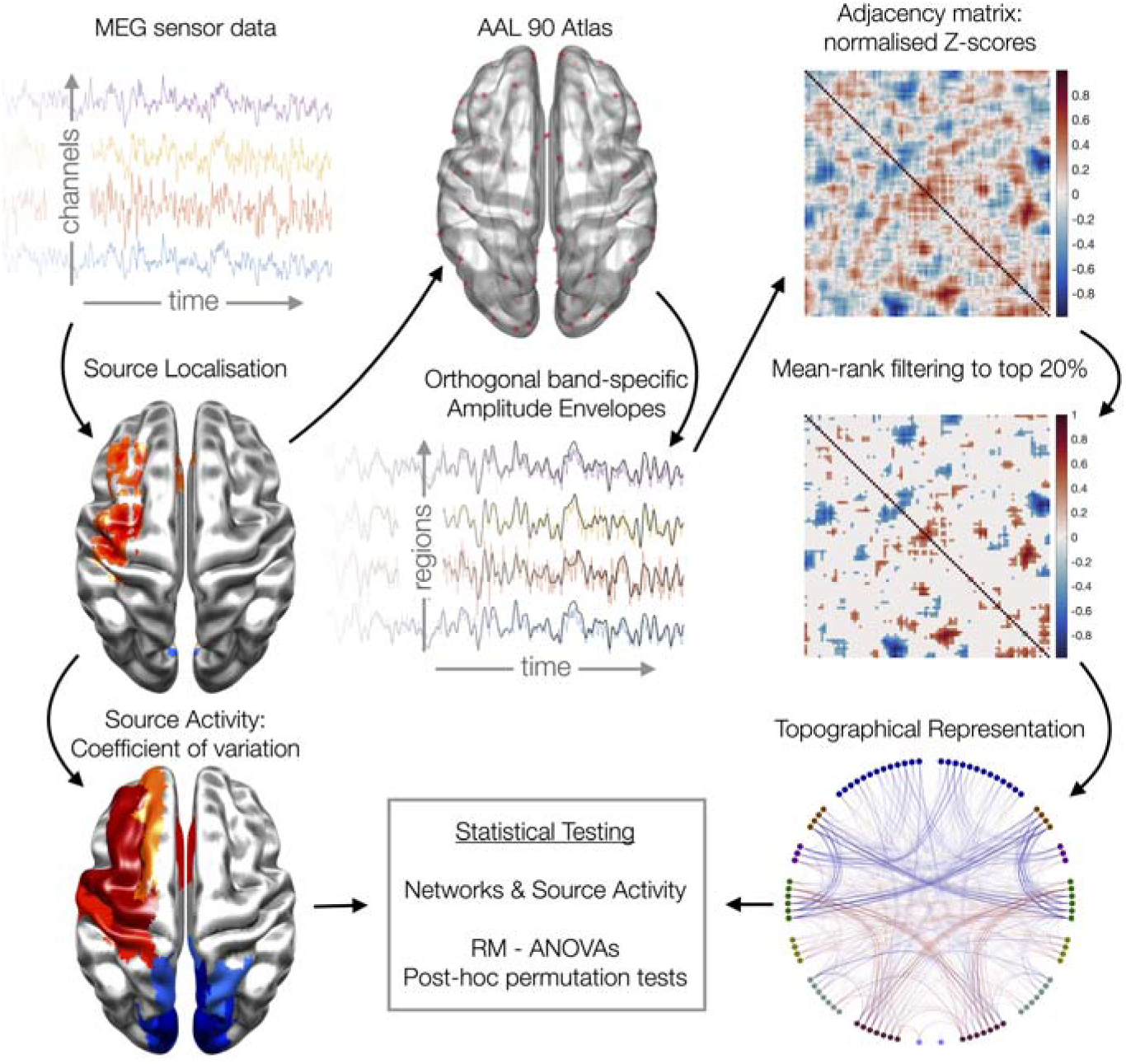
Schematic overview of the analysis and statistical pipeline. RM-ANOVA: repeated measures analysis of variance

Source analysis was performed using the linearly constrained minimum-variance (LCMV) beamformer, a spatial filtering method, as implemented in Fieldtrip (Oostenveld et al., 2011). Using a local-spheres conductivity (forward model). the beamformer estimated the temporal activity of each location of a 6 mm grid (inside the head), as linear combinations of the MEG channels. This step was repeated for each of the frequency bands of interest, with the covariance of the MEG channels being recomputed from appropriately band-pass filtered data. For each of the locations on this 6mm grid, the estimated source timeseries for each trial was then concatenated to form a single timeseries that was then taken forward for both connectivity and activity analysis.

#### Analysis of Amplitude-Amplitude Connectivity

Our analysis proceeded using the methods we previously employed in Koelewijin et al (2019). We first reduced the spatial dimensionality of the source reconstructed data to a set of 90 anatomical loci, described by the Automatic Anatomical Labelling (AAL90) atlas (Tzourio-Mazoyer et al., 2002). We chose one grid source to represent each AAL90 region, selected based on the voxel having the largest temporal standard deviation across the resting-state experiment.

Having reduced to a set of 90 regions, the temporal activity of these sources was orthogonalized with respect to each other region using symmetric orthogonalization (Colclough et al., 2015), which further reduces source leakage. Next, the amplitude envelopes of each region were extracted using the absolute of the (complex) analytical signal derived by Hilbert transform (in MATLAB). This timeseries was then downsampled to a temporal resolution of 1s in order to study connectivity mediated by slow amplitude envelope changes (Brookes et al., 2011). A median filter was used at this point to remove any spikes or transients within the envelope.

The functional connectivity of the 90 regions was then calculated using cross-correlations of the amplitude envelopes of each region. Since correlations are undirected, only the upper triangular portion (minus the diagonals) of the 90-by-90 correlation matrix was computed, requiring 4005 computations for each of the frequency bands tested, for each subject. In order to make these correlations more suitable for statistical analysis, and to correct for the varying length of the final timeseries for each person, each of these correlation coefficients were then transformed to variance-normalised Fisher z-statistics, using a procedure that estimates the effective temporal variance of the node timeseries’ null distribution using surrogates generated by randomisation.

We additionally adjusted each person/session’s connectivity matrix to correct for global session effects (Siems et al., 2016). These effects can be generated by experimental session confounds, effecting SNR, such as head-size, head-motion and position within the MEG helmet. Such correction procedures are common in fMRI connectivity analyses, although there is much debate as to the optimum algorithm to be used for post-hoc standardization (Yan et al., 2013). Here we adopted a variant of z-scoring, in which the null mean and standard-deviation of connectivity, across the matrix, is estimated by fitting a Gaussian (Lowe et al., 1998) to the noise peak (+/- 1SD) of the distribution. This estimated mean and SD is then used to Z-score each Fisher’s Z connectivity value for that person/session.

Finally, in order to only analyse connections which are strongly present in every person/session, we estimated the mean rank of every connection in the matrix and removed the weakest 80%.

#### Source Activity Analysis

For each of the beamformer voxels on the 6×6×6mm reconstruction grid (5061 voxels), we estimated source activity for each voxel, frequency, person and session. Estimating raw power estimates from MEG/EEG data, in source-space, and comparing these across people and sessions is problematic because geometrical effects, which vary from session to session and across the brain, can lead to artefactual inter-session differences (Luckhoo et al., 2014). This is particularly important for beamformer reconstructions in which typical corrections for depth-biases in single-state filter weights can exacerbate these artefactual differences. Here, therefore, we use a normalised measure of “activity”, in which we estimate the amplitude envelope at each of the 5061 locations, using the Hilbert methods described above, and calculate the temporal coefficient of variation (CoV) i.e. the temporal standard deviation of the envelope, divided by the mean of the envelope. Finally, we also z-scored these activity measures using the same gaussian-fit procedure described above for our connectivity measures.

### PET Flumazenil Methods

#### Sample

A group of 16 healthy participants (females: 4, age range: 26-61 years, mean age: 46 years) had been recruited as healthy controls for a larger clinical PET study. No participants included in the study reported a history of neuropsychiatric or neurological conditions or were on any prescribed medications. None of the participants consumed alcohol within 48 hours prior to their PET-FMZ scan. All participants provided written informed consent according to the Declaration of Helsinki. Approvals from the UK Administration of Radioactive Substances Advisory Committee (ARSAC) and the Ethics Committee, Imperial College, Hammersmith Hospital, were obtained. For the creation of the FMZ template, data from six healthy participants with global values close to the mean were selected and raw and right-left reversed (flipped) data were used (12 scans).

#### PET scan parameters and image analysis

All PET data were collected on a 953B Siemens/CTI PET scanner. Dynamic 3D PET data were collected, which consisted of 20 frames over a 90-minute time period. Scans were obtained with axial planes parallel to a horizontal plane passing through the anterior and posterior commissures. The tracer, ∼370 MBq of [^11^C]FMZ, was injected intravenously and arterial blood samples were collected in order to calculate the plasma input function with metabolite correction (Hammers et al., 2003; Lammertsma et al., 1993).

Voxelwise parametric maps of [^11^C]FMZ total volume-of-distribution (V_T_) were calculated from the time-series data and arterial plasma input functions with spectral analysis (Cunningham & Jones, 1993; Lassen et al., 1995), allowing for a blood volume fraction, as detailed previously (Hammers et al., 2008).

#### Template creation

Statistical Parametric Mapping software was used to create the [^11^C]FMZ PET template (SPM99, Wellcome Trust Centre for Human Neuroimaging, University College London, London, UK, https://www.fil.ion.ucl.ac.uk/spm/software/spm99/).

PET data were coregistered to the corresponding MRI data, and left-right-reversed, but not resliced. The MRIs were then normalised to the Montreal Neurological Institute standard space (MNI152). The PETs were spatially normalised using the same transformations as for the MRI images and written out with voxel sizes of 2 mm x 2 mm x 2 mm. Data were then averaged with a softmean function and smoothed with an 8 mm (FWHM) Gaussian kernel to yield the final template.

#### Statistical Analyses

For the connectivity and activity analyses, we employed a repeated-measures analysis of variance (RM-ANOVA), of drug (tiagabine, placebo) by session (pre and 1, 3 and 5 hours post). Where drug x session interaction effects were present, we employed post-hoc paired-*t* testing to explore how effects evolved dynamically across sessions (time).

For the connectivity analysis, which was computed in AAL90 template space for computational tractability, we employed randomisation based post-hoc testing with omnibus-correction (Nichols & Holmes, 2001). For the activity analysis, which was performed at the voxel level (‘voxelwise’), permutation testing was not computationally tractable; as such we instead employed false-discovery-rate (FDR) correction (Benjamini & Hochberg, 1995).

For the comparison of the spatial distribution of activity effects with the PET FMZ V_T_ template, we used randomisation-based Pearson’s correlation (5000 permutations) with omnibus correction.

## Results

### Functional Connectivity

Figure 2 shows connections (edges) demonstrating significant drug-by-session interaction for each frequency band (top row), as well as edges showing a main effect of drug (middle row) and session (bottom row), as computed by RM-ANOVA. Connections showing significant interaction effects were observed in alpha (n=83), beta (62), theta (61) and delta (7) bands, with most connections clustered around posterior regions (occipital / parietal). There were no edges in the gamma range demonstrating an interaction effect (figure 2 demonstrates only gamma^1^, 40 – 60 Hz, however the negative result was also observed in the higher gamma bands). Connections showing a main effect of drug were observed in theta (n=96 edges), alpha (58), beta (33) and delta (20), largely confined to posterior regions but including interactions with cingulate and temporal regions in the alpha band. Effects of session were observed across all frequency bands (n=130 edges in alpha, 111 in beta, 48 in theta, 22 in delta and 3 in gamma^1^). Again, these were posteriorly focussed but included interactions with frontal, deep (delta) and temporal (theta) regions. A list of the top-5 strongest (significant) edges for each frequency band, and each term (effect), are listed in supplementary table 1.

**Figure 2.**
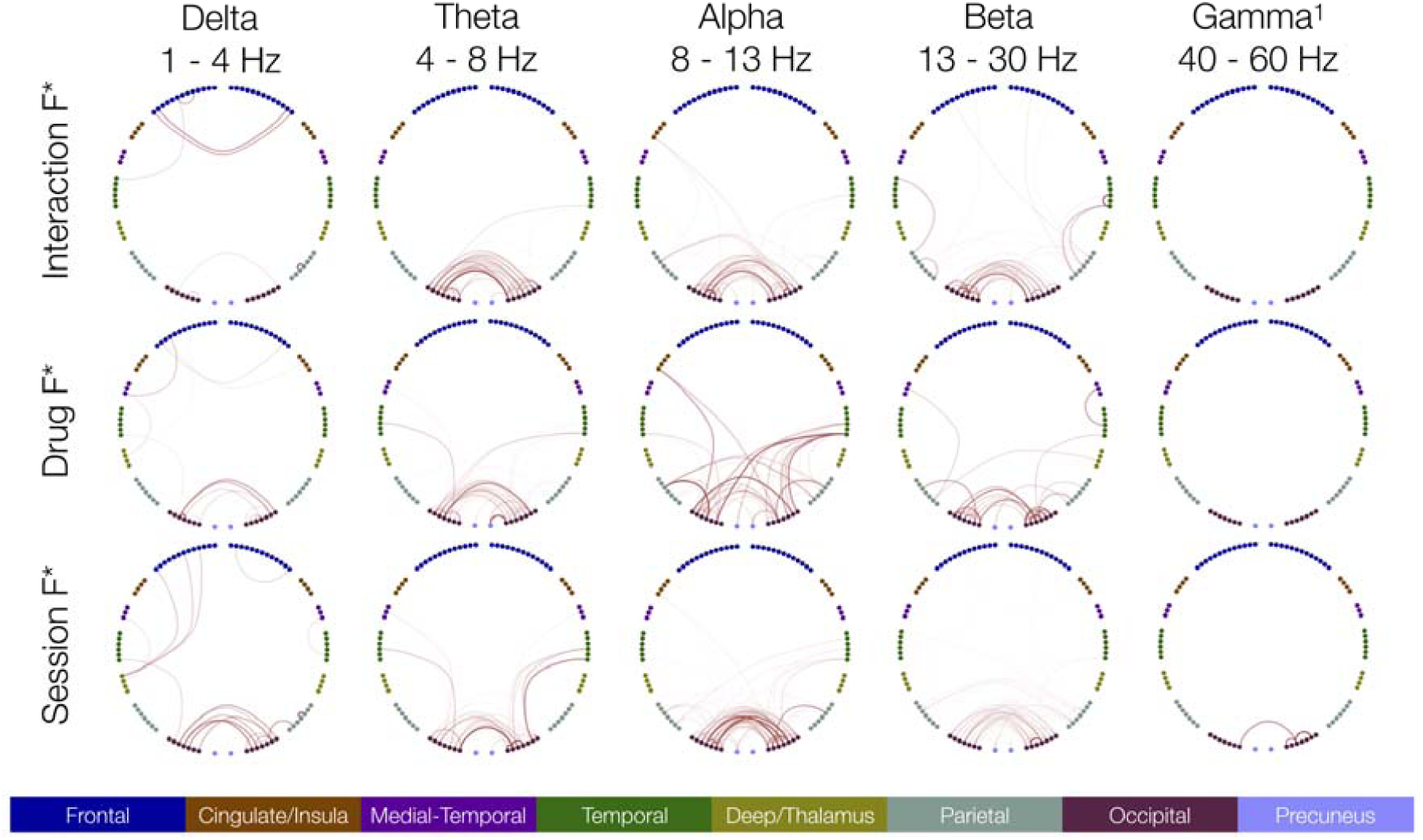
Topographical representation of the significant (p<.05) connections (edges) from the RM-ANOVA, showing interaction effects (top row), as well as drug (row 2) and session effects (row 3). Colour key at bottom describes regions, left/right represents hemisphere (left=left).

*Post-hoc* permutation-based randomisation testing of within-drug session effects revealed significant reductions (figure 3, blue lines) of edge strength for tiagabine at 1, 3 and 5 hr vs. pre, across delta, theta, alpha and beta. Analysis of the topography revealed these reductions were confined to occipital cortices. Increases were observed at two frontal edges (1 at 3 hours, 1 at 5 hours) in the delta band. Significant increases in edge strength were observed between frontal and temporal regions (3 hours vs pre) and temporal and deep regions (5 hours vs pre) across all three gamma windows for placebo. No other within-drug-vs.-pre changes in edge strength were observed for placebo. A list of all significant edges for each frequency band and each comparison (n=6; 1, 3 and 5 hours vs. pre for each drug) are listed in supplementary table 2. Figure 4 summarises the ‘time-course’ of effects on occipital connectivity for each post-drug session vs pre.

**Figure 3.**
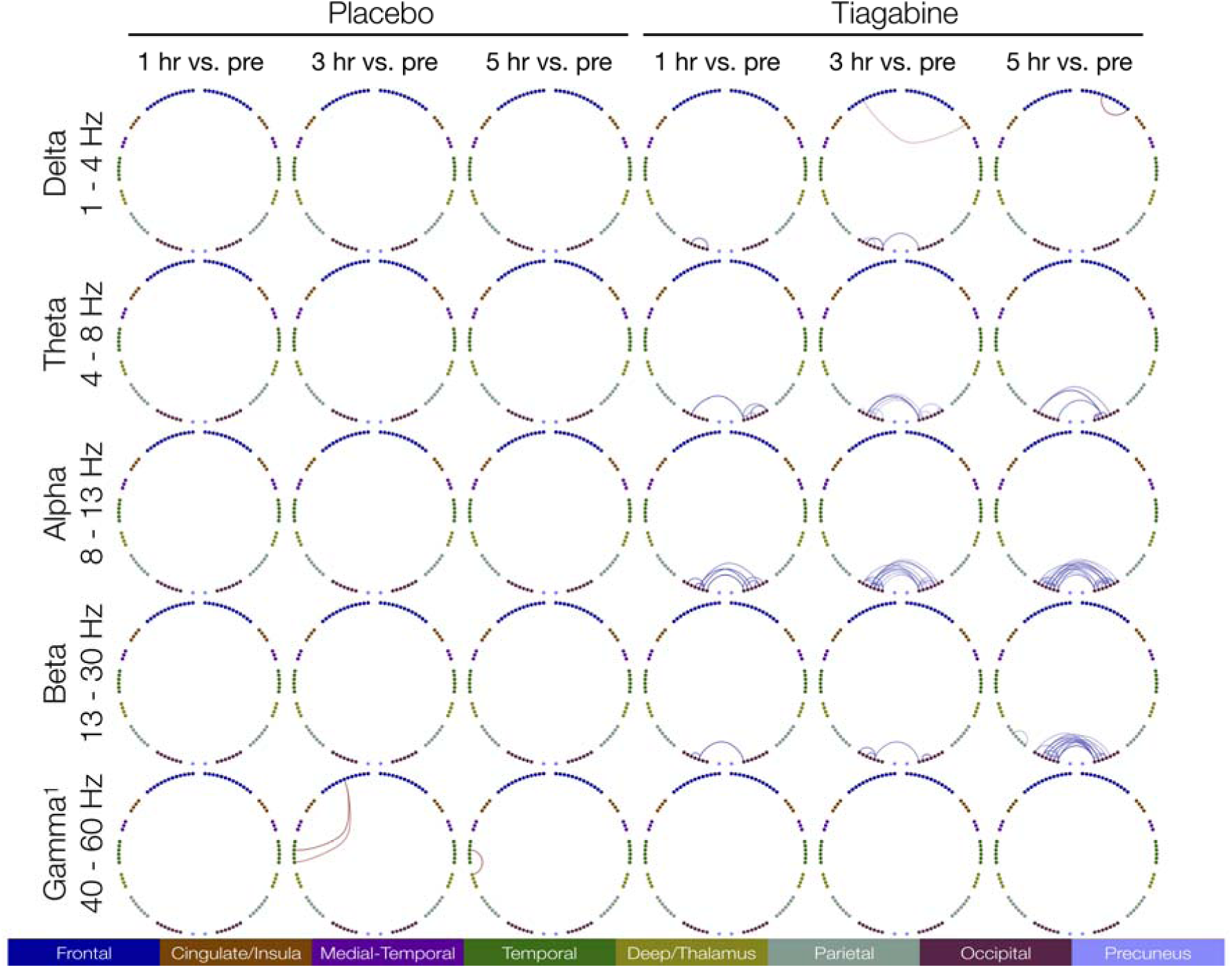
Topographical representation of the post-hoc permutation-based (5k) tests, showing only significant (p<.05) connections, for within-drug comparisons to baseline (1-, 3- and 5-hours vs. pre). Colour represents direction (red = increased vs. pre, blue = decreased vs. pre).

**Figure 4.**
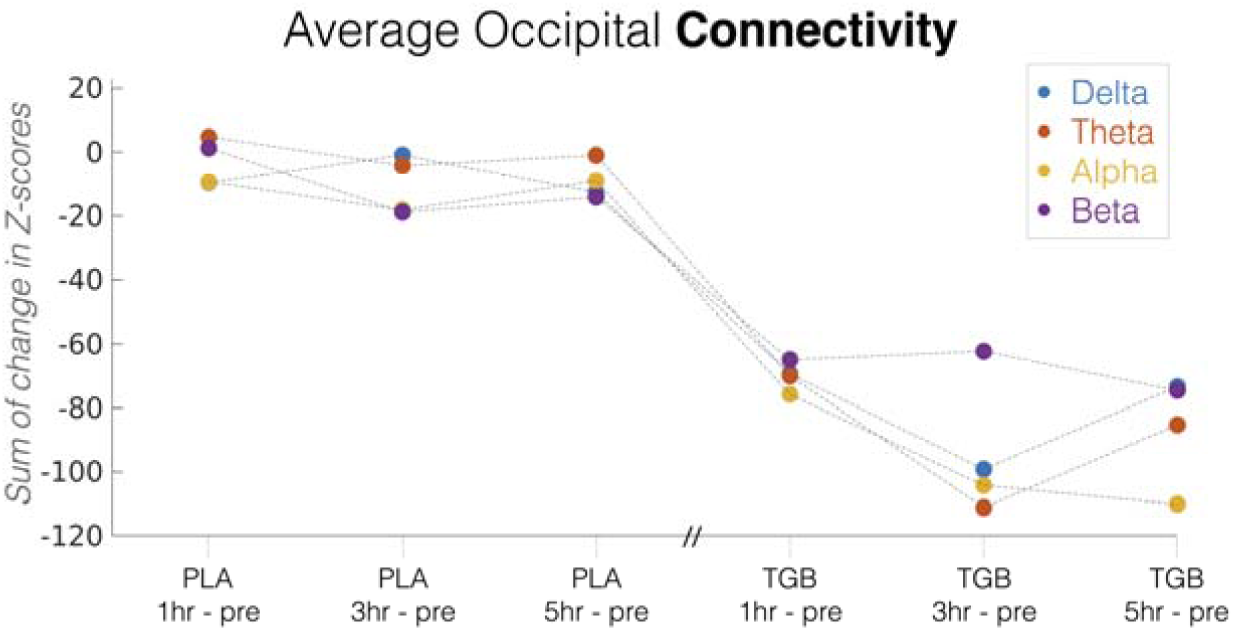
Time course of mean occipital connectivity effects for post-drug vs. baseline, for placebo (PLA) and tiagabine (TGB). Colours represent frequency bands. Note that the Placebo and Tiagabine sessions were performed on different days.

### Source Activity

Figure 5 shows the voxels whose ‘activity’ demonstrated significant drug-by-session interaction, for each frequency band (i.e. voxels with p < 0.05 after false recovery rate correction (FDR)) (Benjamini & Hochberg, 1995). Significant drug x session interaction effects on activity were observed in alpha (n = 3133 voxels), beta (n = 1900), theta (n = 1776) and delta (n = 1539, see figure 6(A)) but not gamma. Within each band, peak F-statistics in size order were F(1,14) = 120.37 (theta), F(1,14) = 106.43 (beta), F(1,14) = 72.87 (delta) and F(1,14) = 67.62 (alpha). These are summarised in figure 6(B).

**Figure 5.**
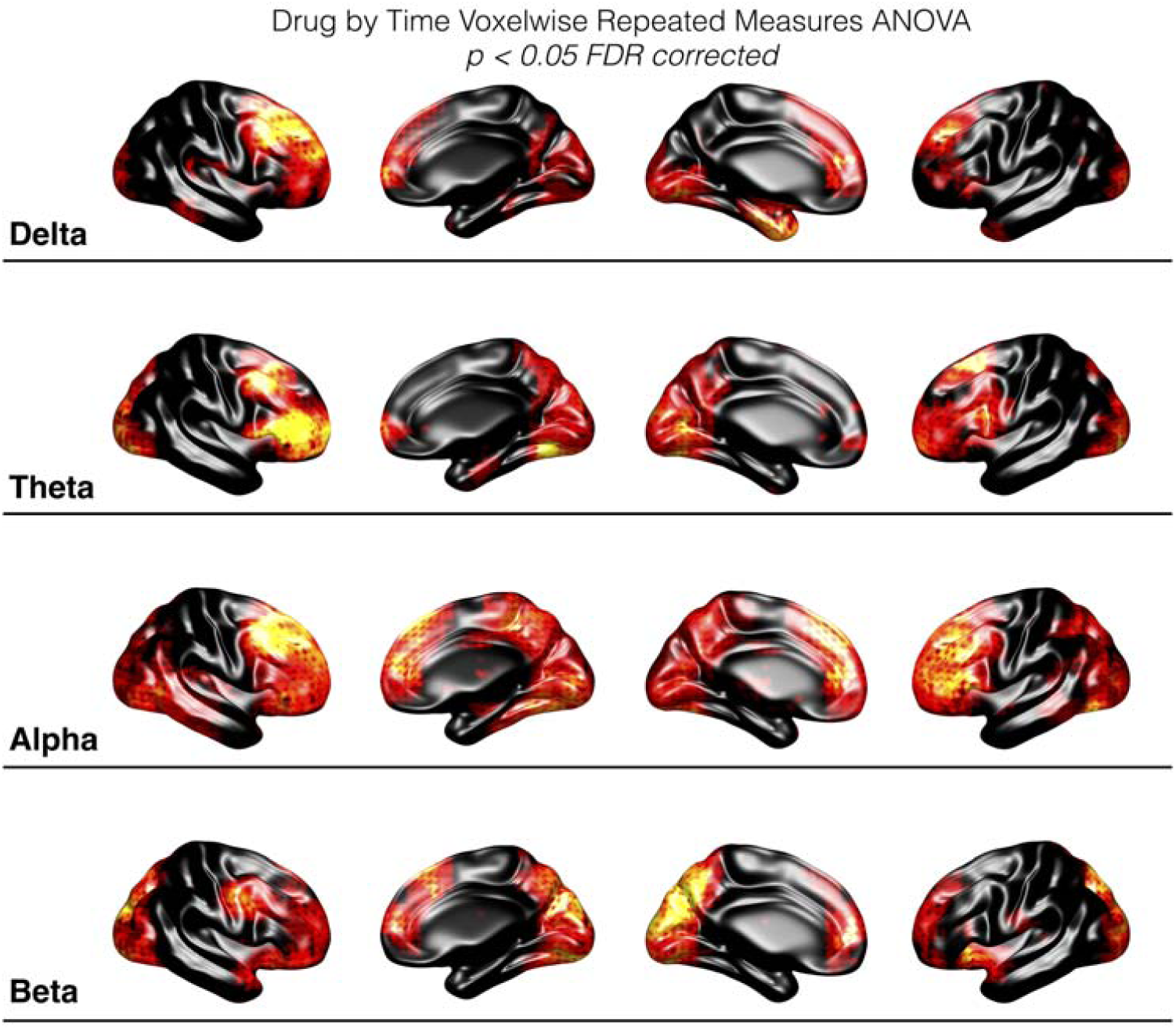
Surface representation of the voxels demonstrating significant (p<.05) drug-by-session interaction effects on source activity (from RM-ANOVA). Red-Yellow colour axis represents relative strength of F-statistic for significant voxels (p<0.05 FDR corrected). Black = n.s.

**Figure 6.**
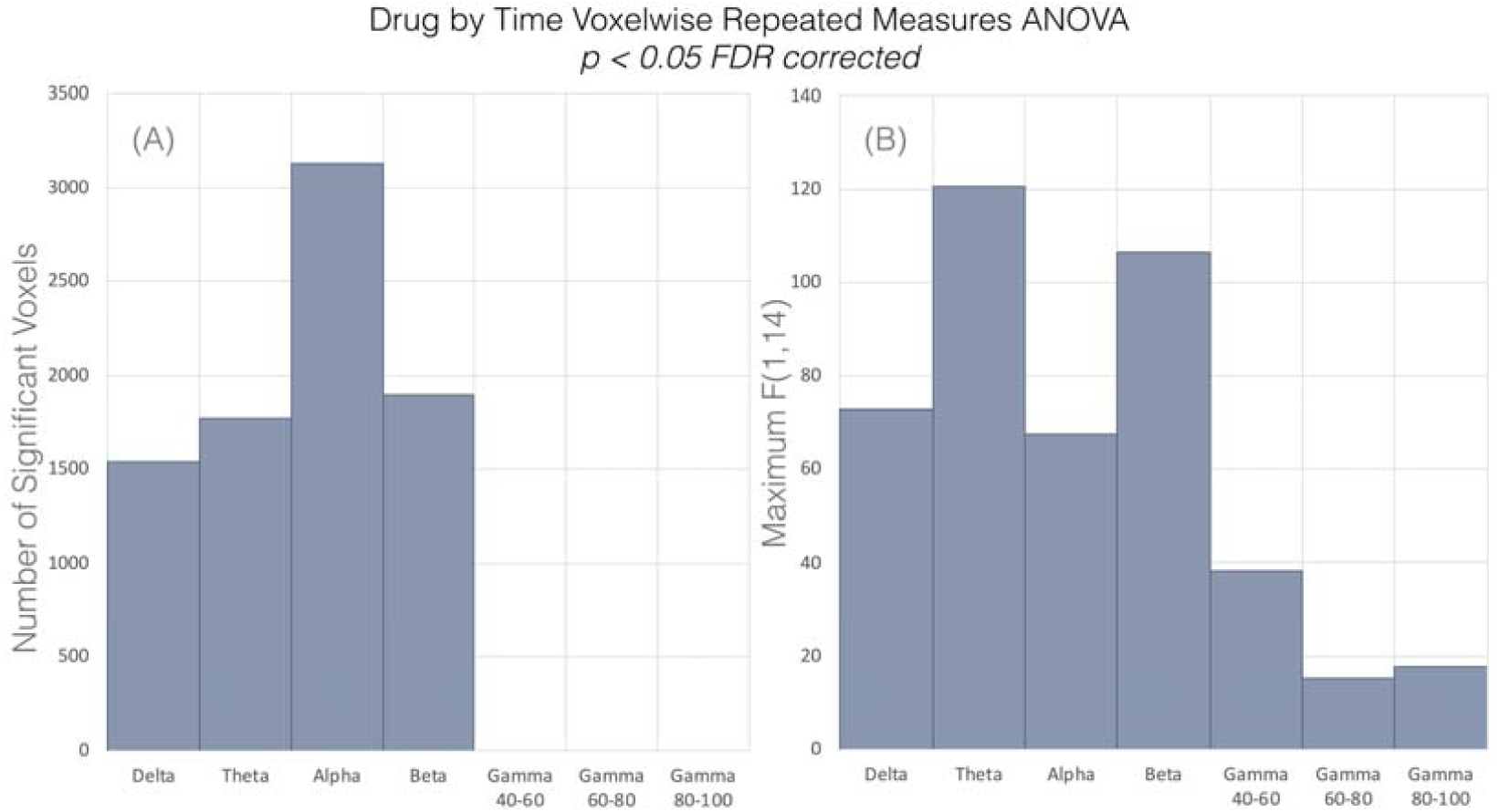
Summary plots for the voxelwise drug-by-session repeated-measures ANOVA, for source activity, showing (A) the number of significant voxels in each frequency band (p < 0.05 FDR corrected) and (B) the maximum F-statistic in each frequency band.

Post-hoc paired-*t* tests between post-drug time points versus the pre-drug recording, were computed for voxels demonstrating significant interaction effects in the repeated measures ANOVA. The resulting *t*-difference maps for each frequency band, shown for tiagabine in figure 7, depict a pattern common across frequencies whereby frontal regions demonstrated increased activity and posterior regions decreased activity, in post-tiagabine sessions relative to pre-drug (this is shown only for tiagabine in figure 7, not placebo). Figure 8 summarises the time-course of these paired-t effects over all sessions (all post-drug vs pre, for both tiagabine and placebo sessions).

**Figure 7.**
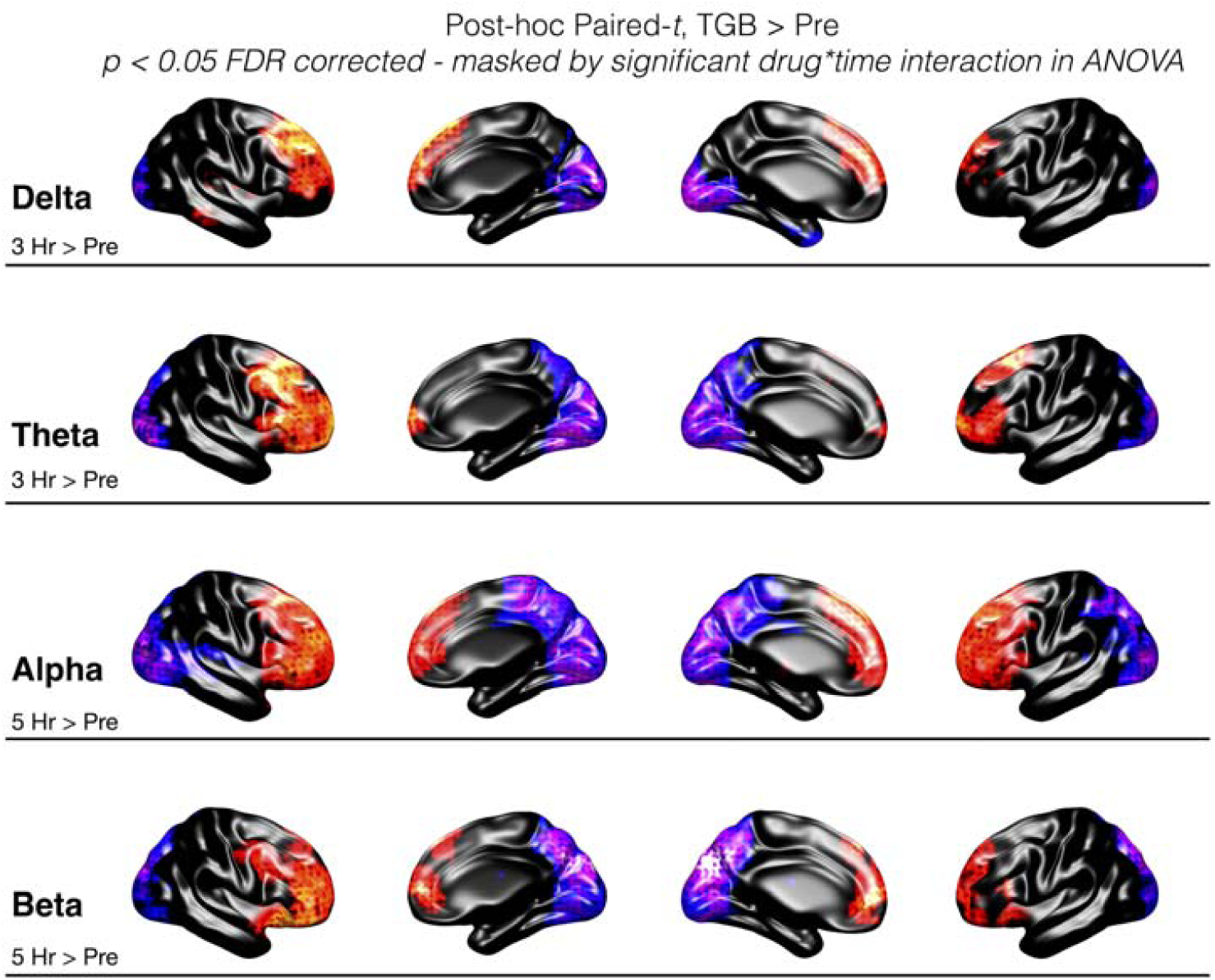
Surface representation of the significant (p < 0.05 FDR corrected) post-hoc paired-t values for post-tiagabine vs pre, for activity measures in each frequency band. The post-drug session depicted was chosen based on the session showing the maximum number of significant voxels (see figure 8). For delta and theta, this was 3 hr post tiagabine vs pre, whereas for alpha and beta it was 5 hr post tiagabine vs pre. Colour axis represents direction of effect, with red/yellow representing significant, positive t-values, blue/purple representing significant negative t-values and black representing non-significant voxels.

**Figure 8.**
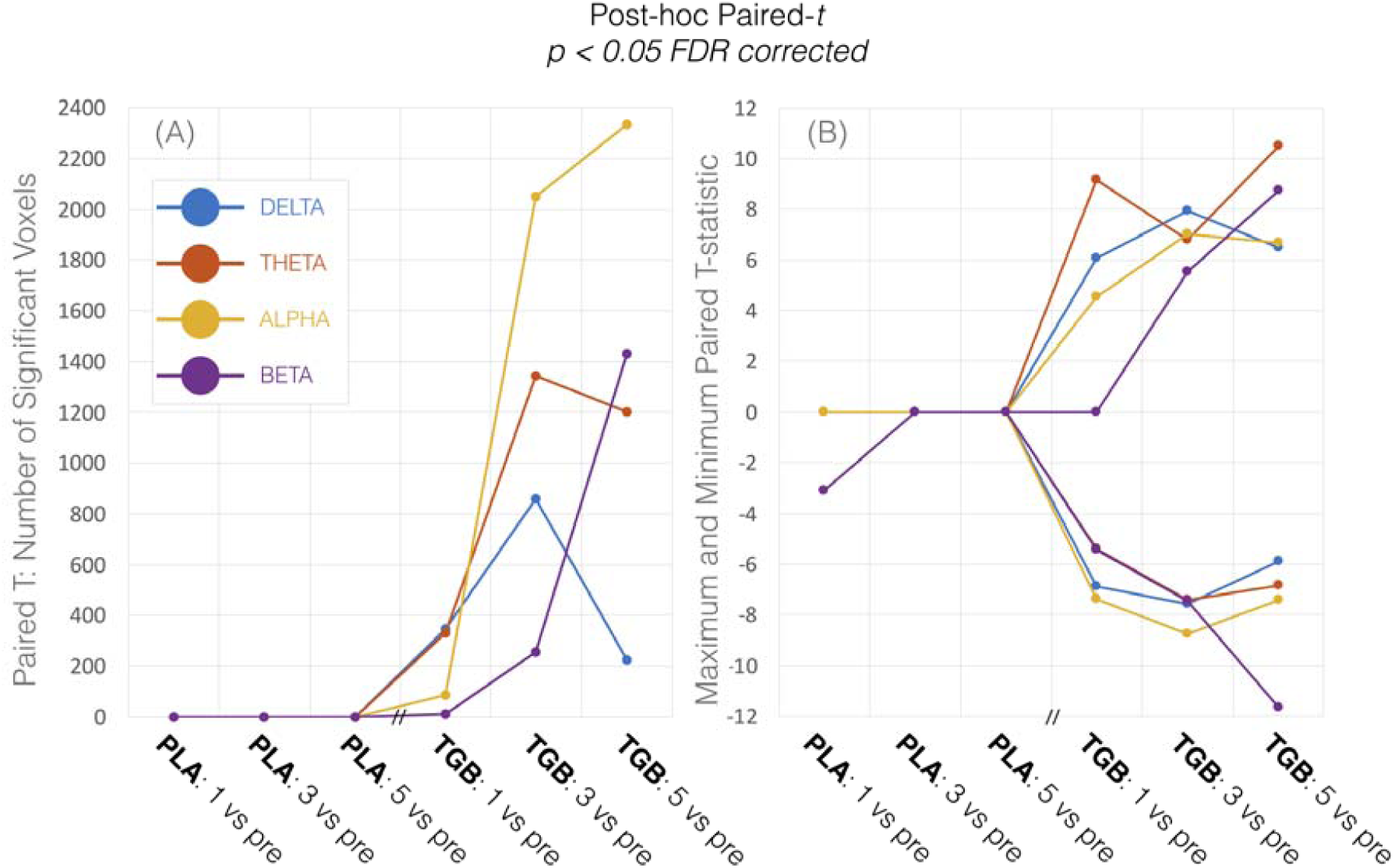
Summary of the source activity post-hoc paired-t effects for post-drug vs pre for each session. (A) shows the number of significant (p < 0.05 FDR corrected) voxels for each frequency band. (B) shows the corresponding peak significant paired t-value for each comparison (x-axis) and frequency band. For clarity, the strongest positive and negative t-values are presented separately. Note that the session demonstrating the peak number of voxels for each frequency band in (A) was selected for graphical representation in figure 7. Note also that the Placebo (PLA) and Tiagabine (TGB) sessions were performed on different days.

### Correlation between the spatial distribution of drug effects and mean Flumazenil PET measures

We observed significant correlations (randomisation testing with 5000 permutations and omnibus correction) in the spatial distribution of the mean drug effects on activity (i.e. absolute paired-*t* value of drug vs pre) with scaled FMZ V_T_, across delta (r=0.15, 0.25 and 0.2 at 1, 3 and 5 hours, respectively), theta (r=0.15, 0.26 and 0.19 at 1, 3 and 5 hours, respectively), alpha (r=.18, 0.22 and 0.24 at 1, 3 and 5 hours, respectively) and beta (r=0.21, 0.23 and 0.24 at 1, 3 and 5 hours, respectively). These correlations suggest that the magnitude of drug effects on activity, irrespective of sign, tended to be bigger in regions with higher FMZ V_T_ (figure 9).

**Figure 9.**
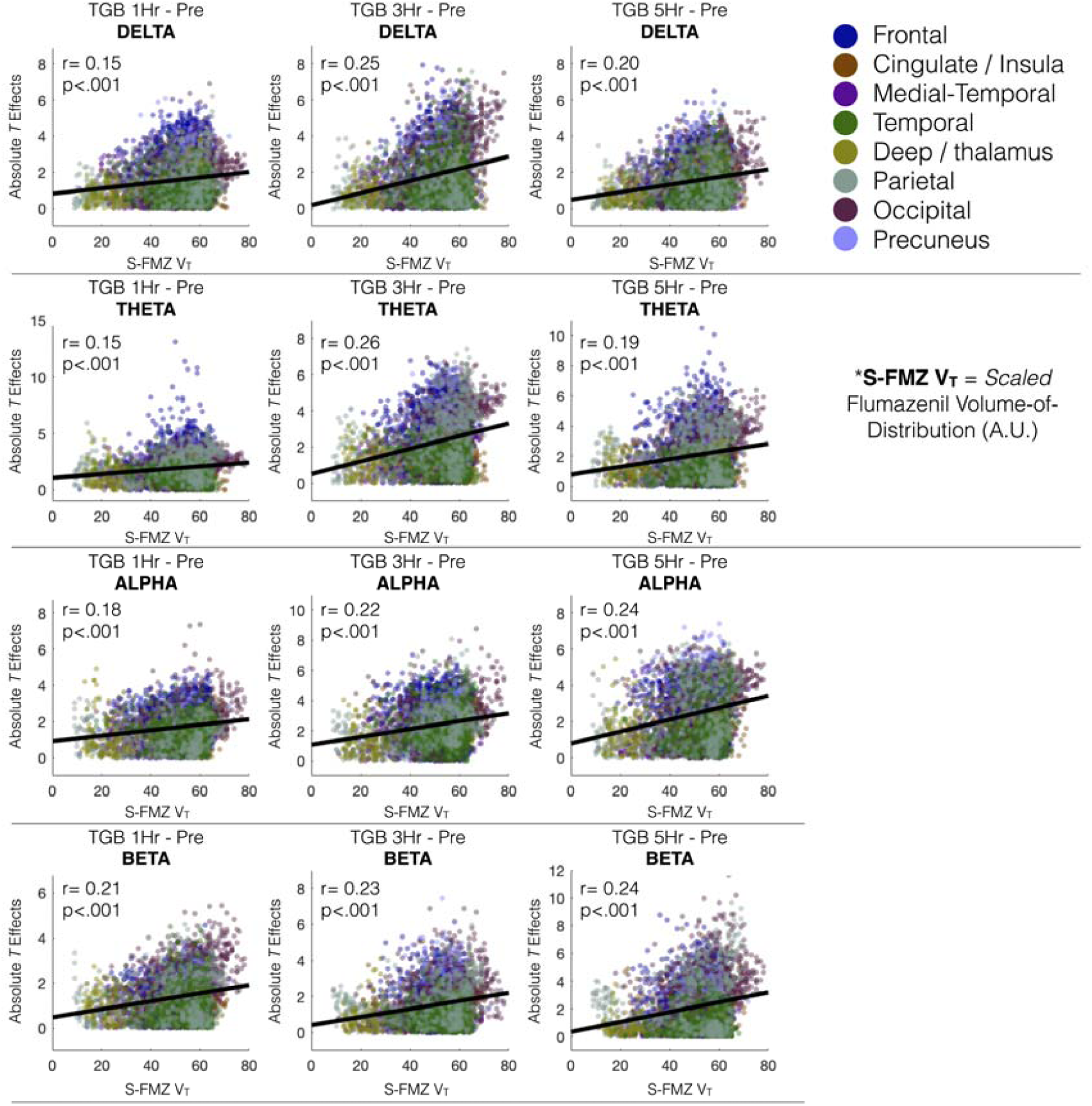
Spatial distribution of scaled flumazenil FMZ V_T_ correlations with effects of drug (absolute *t* value) on activity across frequency bands delta (top), theta, alpha and beta (bottom). Dots represent individual voxels, colour coded by AAL region to which they belong. Statistics (r and p-value) are derived from randomisation-corrected Pearson correlations with omnibus correction for multiple comparisons. Note that because of inherent spatial smoothing within both the PET and MEG beamformer maps, the p-values listed here are likely to be artificially small.

## Discussion

Our results indicate that pharmacologically increased GABA availability leads to reductions of both local activity and regional connectivity across low frequencies from delta, through to higher, beta frequencies. Connectivity effects are localised to posterior brain regions, with predominant effects in occipital lobe. Figure 4 summarises the time course of mean effects for the connectivity, averaged across occipital regions, showing delta, theta, alpha and beta. Voxelwise analysis of ‘activity’ (temporal coefficient of variation) in each frequency band also demonstrated tiagabine induced reductions in posterior regions (figure 7), that were accompanied by increases in activity in frontal regions. Figure 8 summarises the time course of activity effects for each frequency band.

As summarised in figure 10, the effects of tiagabine on functional connectivity across the four frequency bands show a similar temporal pattern, suggesting a common mechanism for the observed effects that is not frequency specific. This wide-band effect is consistent with theories of GABAergic function as a crucial component of oscillation generation, whereby inhibitory GABAergic currents exert temporal control over principal excitatory populations.

**Figure 10.**
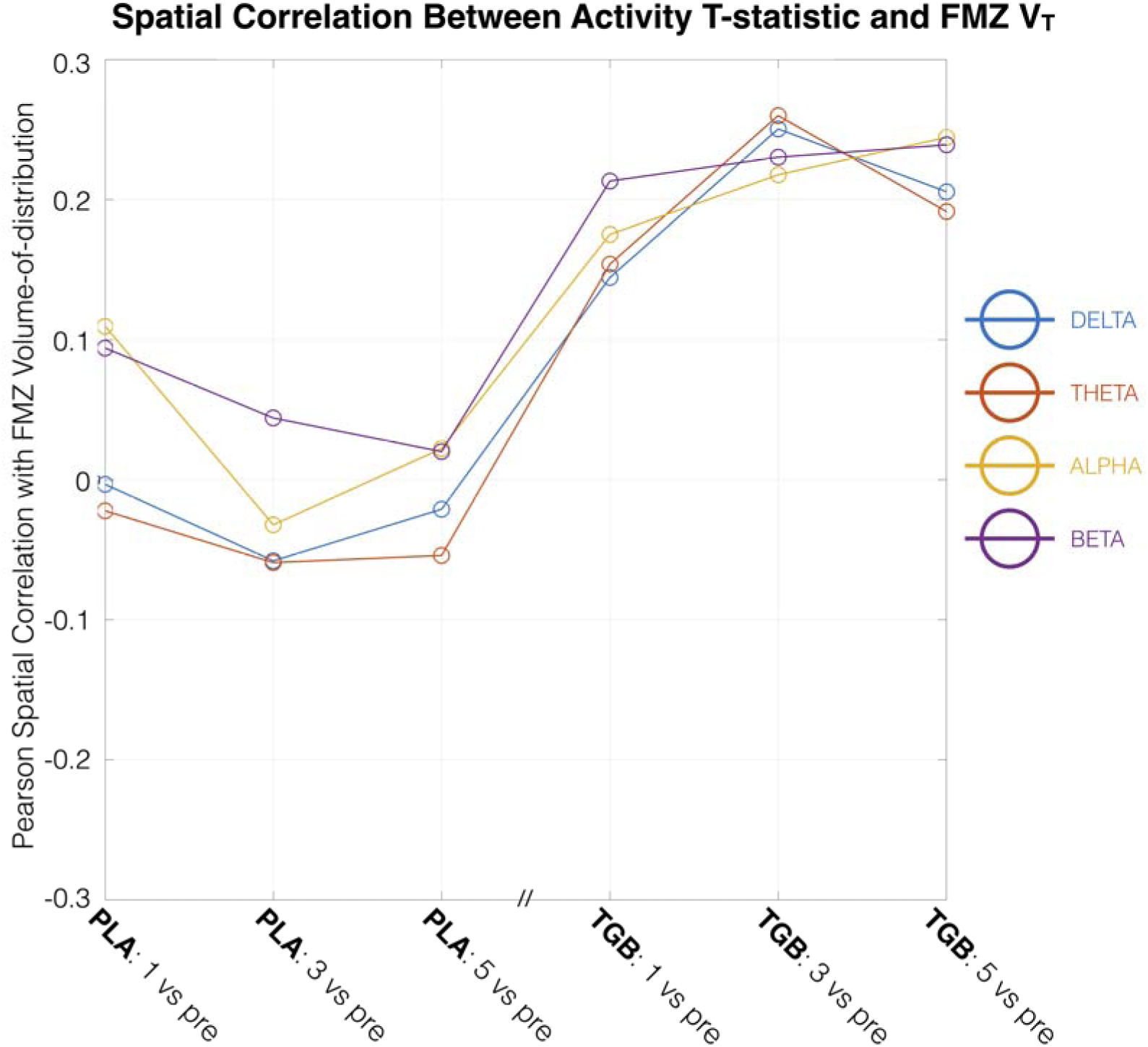
Summarising the spatial correlation of flumazenil volume-of-distribution with drug effects on activity across MEG sessions. Individual correlations for the tiagabine sessions are shown in figure 9. Note that the Placebo (PLA) and Tiagabine (TGB) sessions were performed on different days.

Using a ‘template’ map of flumazenil V_T_, which is a quantitative measure of flumazenil binding at GABA_A_ receptors, we demonstrated that the distribution of drug-induced effects in activity, as measured by the absolute paired-*t* value of post-drug vs pre, correlated with the distribution of GABA_A_ receptors across the brain (figure 9). In other words, regions with higher flumazenil binding also demonstrated bigger effects of drug on activity in delta, theta, alpha and beta bands. The ‘time course’ of this correlation across MEG session is summarised in figure 10.

Previous studies have suggested that tiagabine is not directly active at GABA_A_ sites, instead exerting effects on the GABAergic system through inhibiting reuptake of GABA predominantly through GABA transporter 1 (GAT1) (Madsen et al., 2011; Ransom & Richerson, 2009) and the extrasynaptic betaine/GABA transporter BGT1 (Madsen et al., 2011).

Despite this lack of binding profile at GABA_A_ sites, tiagabine increases the decay time of GABA_A_ mediated inhibitory postsynaptic currents (IPSCs) (Thompson & Gähwiler, 1992) and increases the affinity of GABA_A_ receptors containing the benzodiazepine site to GABA (Frankle et al., 2009). This is important, since it links increased availability of GABA (through inhibited reuptake) to a functional effect or consequence, in the form of increased affinity and IPSC decay time. In other words, it demonstrates that merely increasing the amount of GABA with tiagabine does have a downstream effect on GABAergic function, via a GABA_A_ mediated phenomenon. As such, our findings of tiagabine-attenuated functional connectivity and activity at low frequencies across posterior cortex may reflect enhanced GABAergic function in the form of lengthened IPSCs. The correlation in spatial effects between activity and (template, canonical) FMZ V_T_ supports this, because the effects are spatially colocalised to regions with a higher density of GABA_A_ receptors.

The above argument cannot, however, explain the significant increases in activity observed in frontal regions after administration of tiagabine. Increased frontal power (not activity) of low frequency oscillations has been noted previously, albeit at the sensor level, with the GABA enhancing drugs tiagabine and gaboxodol, but not zolpidem (Nutt et al., 2015).

Inclusion of template flumazenil V_T_ data allowed us to go a step further than quantifying changes in connectivity with enhanced GABA, because we were able to demonstrate evidence that the amount of functional change in a region was predicted by that region’s canonical density of GABA_A_ receptors. This provided a mechanistic link between the drug target and observed response, clearly demonstrating target engagement.

Although we observed connectivity and activity changes across frequency bands ranging from delta to beta, we observed no connectivity changes in the gamma band, even when sub-dividing into discrete 20 Hz windows. One explanation for this, is that the GABAergic processes altered by tiagabine did not disrupt the generative mechanisms of gamma oscillations. However, this is unlikely since the generation of (and spectral characteristics of) gamma oscillations are coupled to GABA_A_ dynamics (Bartos et al., 2007; Whittington et al., 2000). Instead, we propose that gamma oscillations, which are considered locally generated phenomena within cortical columns, are harder to measure between-region. This is because gamma oscillations, by virtue of being fast, can exist more transiently than lower frequencies, and are therefore harder to detect when correlating long amplitude envelopes. If this is the case, methods aimed at identifying fast dynamical switching, such as Hidden Markov Models, may be more successful for quantifying faster frequencies (Baker et al., 2014).

### Limitations

This study used a template of flumazenil V_T_ derived from existing data, and from different subjects to those who took part in the tiagabine MEG study. Using averaged PET data as a canonical reference map of GABA_A_ density across the brain permitted us to compare to the average distribution of drug effects across the brain, but lacks the specificity afforded by having PET and MEG on the same individuals. Future studies should consider this, since having both would allow for detailed analysis of individual differences. A caveat here is the relative expense and exposure to radioactivity with PET compared with the safety of M/EEG.

Our investigation of the spatial co-distribution of activity with FMZ V_T_ may overestimate correlations due to the inherent smoothness in the PET images. This smoothness-induced autocorrelation reduces the effective degrees of freedom, thereby rendering inflated statistics in conventional correlation measures, such as the Pearson’s correlation employed here (Afyouni et al., 2019; Arbabshirani et al., 2014).

### Conclusions

We have demonstrated that pharmacologically increased GABA availability led to increased frontal activity and reduced posterior activity – and inter-region connectivity – across multiple frequency bands. The spatially distinct pattern of activity changes correlated with the distribution of GABA_A_ receptors across the brain. We propose a mechanistic explanation for our results, whereby increased GABA availability (by tiagabine) led to increased affinity of GABA_A_ receptors for GABA (Frankle et al., 2009), lengthening IPSCs (Thompson & Gähwiler, 1992) and consequently reducing low frequency oscillatory activity and connectivity (by the lengthened IPSCs ‘dampening’ current fluctuations underpinning macroscopic oscillations). While demonstrating a role for GABAergic dynamics in shaping the activity and topography of functional connectivity, we have also demonstrated the utility of MEG-based measures of connectivity and activity as a tool for time and frequency-resolved electrophysiological receptor mapping.

## Acknowledgements

The work presented here was supported by CUBRIC and the School of Psychology at Cardiff University as well as an MRC UK MEG Partnership Grant, MR/K005464/1. ADS & HLC are supported by a Wellcome Strategic Award (104943/Z/14/Z).

AH acknowledges support by the Wellcome EPSRC Centre for Medical Engineering at King’s College London (WT 203148/Z/16/Z) and the Department of Health via the National Institute for Health Research (NIHR) comprehensive Biomedical Research Centre award to Guy’s & St Thomas’ NHS Foundation Trust in partnership with King’s College London and King’s College Hospital NHS Foundation Trust. Data for the FMZ template were originally acquired at the MRC Clinical Sciences Centre, Hammersmith Hospital.

The brain surfaces in this paper were rendered using the SourceMesh toolbox (https://github.com/alexandershaw4/SourceMesh).

