## supplementary for "GABA_A_ receptor mapping in human using non-invasive electrophysiology"

Supplementary Tables

Supplementary Table 1:

Functional Connectivity rm-ANOVA, top 5 most significant connections for each frequency band.

| **Delta (1 – 4 Hz)** | | | | |
| --- | --- | --- | --- | --- |
| Test | Region 1: | Region 2: | F | p |
| Interaction | Parietal Inf R | SupraMarginal R | 6.0174 | 0.0009 |
|  | Frontal Med Orb L | Frontal Med Orb R | 4.9490 | 0.0033 |
|  | Rectus L | Rectus R | 4.6599 | 0.0046 |
|  | Frontal Inf Oper L | Rolandic Oper L | 4.4515 | 0.0059 |
|  | Calcarine L | Lingual L | 3.6115 | 0.0166 |
| Drug | Occipital Sup L | Occipital Mid L | 13.5278 | 0.0009 |
|  | Occipital Mid L | Fusiform R | 10.7155 | 0.0028 |
|  | Occipital Mid L | Occipital Inf R | 10.3599 | 0.0032 |
|  | Olfactory L | Amygdala L | 8.7925 | 0.0061 |
|  | Lingual R | Occipital Sup R | 8.6972 | 0.0063 |
| Session(Time) | Parietal Sup R | Parietal Inf R | 5.8773 | 0.0010 |
|  | Cuneus R | Occipital Sup L | 5.0906 | 0.0027 |
|  | Lingual L | Lingual R | 4.8874 | 0.0035 |
|  | Occipital Inf R | Angular R | 4.7116 | 0.0043 |
|  | Occipital Mid L | Fusiform R | 4.5632 | 0.0051 |
| **Theta (4 – 8 Hz)** | | | | |
| Interaction | Calcarine R | Occipital Sup L | 9.6813 | 1.5e-05 |
|  | Calcarine L | Occipital Sup L | 7.9987 | 9.4e-05 |
|  | Lingual R | Fusiform R | 7.2188 | 0.0002 |
|  | Calcarine R | Occipital Mid L | 6.9252 | 0.0003 |
|  | Occipital Sup R | Occipital Inf R | 6.7322 | 0.0003 |
| Drug | Calcarine R | Precuneus R | 27.5428 | 1.4e-05 |
|  | Occipital Sup L | Occipital Inf R | 23.6887 | 3.9e-05 |
|  | Occipital Sup L | Occipital Inf L | 20.3373 | 0.0001 |
|  | Calcarine L | Occipital Sup L | 20.3050 | 0.0001 |
|  | Lingual R | Occipital Sup L | 19.9387 | 0.0001 |
| Session(Time) | Calcarine R | Cuneus R | 9.1714 | 2.5e-05 |
|  | Calcarine L | Cuneus R | 9.0380 | 2.9e-05 |
|  | Cuneus R | Occipital Sup R | 8.7934 | 3.8e-05 |
|  | Occipital Inf R | Temporal Inf R | 7.8617 | 0.0001 |
|  | Calcarine L | Fusiform L | 6.4115 | 0.0005 |

| **Alpha (8 – 13 Hz)** | | | | |
| --- | --- | --- | --- | --- |
| Test | Region 1: | Region 2: | F | p |
| Interaction | Occipital Sup R | Occipital Mid R | 10.2516 | 8e-06 |
|  | Cuneus L | Occipital Sup L | 9.9289 | 1.1e-05 |
|  | Cuneus R | Lingual L | 9.4630 | 1.8e-05 |
|  | Cuneus L | Lingual L | 8.9448 | 3.3e-05 |
|  | Occipital Sup L | Occipital Mid R | 8.4851 | 5.5e-05 |
| Drug | Occipital Sup L | Parietal Sup L | 12.4522 | 0.0014 |
|  | Cuneus L | Temporal Inf R | 11.6871 | 0.0019 |
|  | Occipital Inf R | Fusiform L | 11.5644 | 0.0020 |
|  | Fusiform R | Temporal Inf R | 11.0559 | 0.0024 |
|  | Lingual R | Parietal Sup R | 10.2943 | 0.0033 |
| Session /Time | Lingual R | Occipital Sup L | 14.2847 | 1e-07 |
|  | Cuneus L | Occipital Mid R | 13.2947 | 3e-07 |
|  | Cuneus L | Occipital Sup R | 13.1584 | 4e-07 |
|  | Cuneus L | Lingual L | 12.4191 | 8e-07 |
|  | Cuneus L | Occipital Sup L | 12.2823 | 9e-07 |
| **Beta (13 – 30 Hz)** | | | | |
| Interaction | Lingual L | Occipital Mid L | 8.4878 | 5.5e-05 |
|  | Temporal Mid R | Temporal Inf R | 8.2960 | 6.8e-05 |
|  | Calcarine R | Lingual L | 7.6925 | 0.0001 |
|  | Lingual L | Occipital Sup L | 7.5542 | 0.0002 |
|  | Calcarine L | Lingual L | 6.7530 | 0.0004 |
| Drug | Cuneus R | Occipital Sup R | 7.7053 | 0.0097 |
|  | Occipital Mid L | Occipital Inf L | 7.3692 | 0.0112 |
|  | Calcarine R | Occipital Mid R | 7.0968 | 0.0126 |
|  | Lingual R | Occipital Inf R | 6.7618 | 0.0147 |
|  | Lingual R | Occipital Mid L | 6.6841 | 0.0152 |
| Session /Time | Cuneus R | Occipital Sup R | 13.1463 | 4e-07 |
|  | Occipital Sup L | Occipital Sup R | 8.6748 | 4.4e-05 |
|  | Occipital Sup R | Occipital Mid R | 8.3095 | 6.6e-05 |
|  | Postcentral L | Parietal Sup L | 6.8519 | 0.0003 |
|  | Cuneus R | Occipital Sup L | 6.8477 | 0.0003 |

Supplementary Table 2:

Post-hoc connectivity, permutation significant connections (Tgb = tiagabine, Pla = placebo):

| **Frequency Band** | **Session [Time]** | **Region 1** | **Region 2** | **Permutation significant paired-*t*** |
| --- | --- | --- | --- | --- |
| Delta | Tgb 1hr v Pre | Occipital Mid L | Calcarine L | -3.3557 |
| Delta | Tgb 3hr v Pre | Occipital Mid L | Calcarine L | -3.8047 |
| Delta | Tgb 3hr v Pre | Calcarine R | Calcarine L | -3.5346 |
| Delta | Tgb 3hr v Pre | Cingulum Mid R | Sup MotorArea L | 2.9982 |
| Delta | Tgb 3hr v Pre | Occipital Inf L | Lingual L | -2.9078 |
| Delta | Tgb 5hr v Pre | Rectus R | Frontal Inf Tri R | 3.0863 |
| Theta | Tgb 1hr v Pre | Occipital Mid L | Calcarine R | -4.1481 |
| Theta | Tgb 1hr v Pre | Fusiform R | Lingual R | -3.9182 |
| Theta | Tgb 1hr v Pre | Fusiform R | Calcarine R | -3.6981 |
| Theta | Tgb 1hr v Pre | Cuneus R | Calcarine R | -3.6067 |
| Theta | Tgb 1hr v Pre | Occipital Sup R | Cuneus R | -3.4947 |
| Theta | Tgb 3hr v Pre | Occipital Mid L | Calcarine R | -5.5462 |
| Theta | Tgb 3hr v Pre | Occipital Mid L | Lingual L | -4.5941 |
| Theta | Tgb 3hr v Pre | Cuneus R | Calcarine R | -4.4740 |
| Theta | Tgb 3hr v Pre | Fusiform R | Occipital Inf R | -4.4594 |
| Theta | Tgb 3hr v Pre | Occipital Sup L | Calcarine R | -4.4368 |
| Theta | Tgb 5hr v Pre | Occipital Inf L | Lingual R | -4.1205 |
| Theta | Tgb 5hr v Pre | Cuneus R | Calcarine R | -4.1185 |
| Theta | Tgb 5hr v Pre | Occipital Sup R | Cuneus R | -4.0029 |
| Theta | Tgb 5hr v Pre | Lingual R | Calcarine R | -3.9878 |
| Theta | Tgb 5hr v Pre | Lingual R | Calcarine L | -3.9037 |
| Beta | Tgb 1hr v Pre | Calcarine R | Calcarine L | -3.4467 |
| Beta | Tgb 1hr v Pre | Occipital Sup L | Cuneus L | -3.40239 |
| Beta | Tgb 1hr v Pre | Lingual R | Cuneus L | -3.3576 |
| Beta | Tgb 1hr v Pre | Lingual L | Cuneus R | -3.3426 |
| Beta | Tgb 1hr v Pre | Occipital Inf L | Cuneus L | -3.2903 |
| Beta | Tgb 3hr v Pre | Occipital Sup L | Cuneus L | -5.2349 |
| Beta | Tgb 3hr v Pre | Occipital Mid R | Occipital Sup L | -4.8263 |
| Beta | Tgb 3hr v Pre | Occipital Inf L | Calcarine L | -4.6820 |
| Beta | Tgb 3hr v Pre | Occipital Sup L | Lingual L | -4.5291 |
| Beta | Tgb 3hr v Pre | Occipital Sup L | Calcarine L | -4.4538 |
| Beta | Tgb 5hr v Pre | Occipital Mid R | Occipital Sup R | -5.3244 |
| Beta | Tgb 5hr v Pre | Occipital Sup L | Cuneus L | -5.1661 |
| Beta | Tgb 5hr v Pre | Occipital Mid R | Lingual L | -4.8676 |
| Beta | Tgb 5hr v Pre | Occipital Inf L | Calcarine L | -4.5433 |
| Beta | Tgb 5hr v Pre | Occipital Mid R | Cuneus R | -4.4988 |
| Alpha | Tgb 1hr v Pre | Lingual L | Calcarine R | -3.2262 |
| Alpha | Tgb 1hr v Pre | Occipital Mid L | Lingual L | -3.0326 |
| Alpha | Tgb 3hr v Pre | Occipital Sup R | Cuneus R | -3.7912 |
| Alpha | Tgb 3hr v Pre | Occipital Mid L | Lingual L | -3.5738 |
| Alpha | Tgb 3hr v Pre | Fusiform L | Lingual L | -3.5514 |
| Alpha | Tgb 3hr v Pre | Lingual L | Calcarine R | -3.4420 |
| Alpha | Tgb 5hr v Pre | Calcarine R | Calcarine L | -4.1660 |
| Alpha | Tgb 5hr v Pre | Lingual L | Calcarine R | -3.9315 |
| Alpha | Tgb 5hr v Pre | Occipital Sup R | Cuneus R | -3.8272 |
| Alpha | Tgb 5hr v Pre | Occipital Mid L | Lingual L | -3.8259 |
| Alpha | Tgb 5hr v Pre | Occipital Mid L | Calcarine L | -3.7461 |
| Gamma | Pla 3hr v Pre | Temp Pole Sup L | Frontal Inf Orb L | 2.9082 |
| Gamma | Pla 3hr v Pre | Temporal Inf L | Frontal Inf Orb L | 2.7081 |
| Gamma | Pla 5hr v Pre | Temp Pole Sup L | Caudate L | 2.8408 |
